# Loss of a major venom toxin gene in a Western Diamondback rattlesnake population

**DOI:** 10.1101/2025.02.02.636150

**Authors:** Noah L. Dowell, Elizabeth Cahill, Sean B. Carroll

**Author notes:** Correspondence and request for materials should be addressed to Sean B. Carroll. Competing interest statement: The authors declare no competing interests.

## Abstract

The biochemical complexity and evolutionary diversity of snake venom composition reflects adaptation to the diversity of prey in their diets. However, the genetic mechanisms underlying the evolutionary diversity of venoms are not well understood. Here, we explored the potential extent of and genetic basis for venom protein variation in the widely-distributed Western Diamondback rattlesnake (*Crotalus atrox*). As in many rattlesnake venoms, metalloproteinases (SVMPs) are the major component of *C. atrox* venom, with three proteins belonging to three distinct major structural SVMP classes, MDC4, MAD3a, and MPO1, constituting the most abundant SVMPs. We found that while most venom proteins, including MDC4 and MAD3a, vary little among individuals, the MPO1 protein is completely absent from some animals, most commonly those from the western part of the species’ geographic range. This distribution correlates with the previous finding of two distinct lineages within *C. atrox* and indicates that different ecological factors have shaped venom composition across the species’ range. We further show that the loss of MPO1 expression is not due to transcriptional down-regulation, but to several independent inactivating mutations at the locus, including whole gene deletion. The recurrent inactivation of a major toxin gene within a *C. atrox* population may reflect relaxed selection on the maintenance of *MPO1* function, but we also raise the possibility that the loss of venom components may be favored if there is a cost to producing a less effective toxin in protein-rich venoms.

## Introduction

Key ecological traits often evolve through co-evolutionary interactions among organisms (e.g. predator-prey, plant-herbivore, host-parasite) [1–3]. Such interactions can drive trait diversification and may ultimately facilitate the exploitation of new environments and resources. A major priority in evolutionary biology is to identify the genes underlying key traits and the molecular basis of functional variation within them.

Animal venoms are key ecological traits that are most often employed for subduing prey and venom composition coevolves within the framework of predator-prey interactions [4–11]. Snake venoms consist of protein mixtures that often vary in composition and sometimes mode of action even between closely related species [12–14]. One general question concerning the molecular evolution of venom composition is the extent to which differences between species are due to differences in gene content or gene regulation. In rattlesnakes, one of the more studied groups, there is evidence for gene content differences underlying venom divergence including both the expansion of certain toxin gene families [15] and the loss of gene family members in certain lineages [16,17].

Of course, such interspecific differences must initially arise through variation within species, and there is growing evidence from field studies of dynamic variation associated with both geography and prey [18–23]. For example, in the widely distributed rattlesnake *Crotalus viridis viridis*, variation in venom composition occurs along the north-south axis of its range and is associated with differences in the availability of different types of prey [24]. Similarly, the discovery of ground squirrels resistant to the effects of sympatric *C. oreganus oreganus* rattlesnake venom indicates an ongoing evolutionary arms race in which both venom toxicity and prey resistance evolve adaptively in response to each other [6]. In *C. adamanteus*, both adult and juvenile snakes from discrete geographic populations express venoms with variable composition [25,26]. While the specific genetic bases of venom variation within these species are not yet known, it has been shown in other rattlesnakes that polymorphisms in venom type (neurotoxic v. hemorrhagic) within several species are due to differences in gene content [17,22]. The discovery of gene loss underlying venom diversity and variation suggests that venom components may be dispensable under some conditions and that the evolution of venom diversity does not occur solely through the addition of new toxins.

Here, to further investigate the extent and potential genetic basis of variation in snake venoms, we have explored variation in *C. atrox* because this species occupies a wide expanse and range of habitats in North America and possesses the largest known venom toxin gene family with at least 30 members of the snake venom metalloproteinase family [15]. Venom MPs are the most abundant component of *C. atrox* venom (by weight) and are primarily responsible for its hemorrhagic activity [27]. Three structural classes (P-I, P-II and P-III) of venom MPs are expressed by distinct genes that encode different combinations of the class-defining domains (e.g. metalloproteinase; metalloproteinase and disintegrin; and metalloproteinase, disintegrin and cysteine-rich for classes P-I, P-II, and P-III, respectively) [15,28]. All three classes of MP can damage micro vessels and contribute to hemorrhage but the extent of damage and mode of action varies between the MP classes [29].

We find that most venom proteins vary little in expression among individuals including the abundantly expressed metalloproteinases (MPs) MDC4 and MAD3a. However, MPO1, the only P-I class MP in the species, is abundant in some specimens but absent in others due to a variety of gene-inactivating mutations (substitutions and gene deletions). Furthermore, the loss of MPO1 expression occurs predominantly in animals found in the western part of the species range, indicating that there are different requirements for venom composition and MPO1 function across the species range. We suggest that the loss of venom components within a population may be due to relaxation of selection and neutral drift but could be favored if there is a cost to producing a less effective toxin.

## Methods

### Mass spectrophotometry of total venom

Sample preparation and mass spectrophotometry of *C. atrox* venoms was performed as described previously [30]. The high sequence similarity between venom metalloproteinases prompted us to use counts of exclusive unique peptides (or spectra) to assess venom protein abundance. Additionally, percent peptide coverage of the metalloproteinase domain was used to distinguish between expression of a complete or partial protein. The detection of any peptides from a specimen that is inferred to have deleted that gene could be due to technical reasons, for example, low level instrument contamination between specimens or perhaps miss-assignment of a particular peptide to a specific protein due to conserved sequences. Using exclusive unique peptides and coverage can minimize but does not eliminate challenges surrounding accurate determination of which proteins in large gene families are present/absent from venom.

### Venom gland RNA sequencing libraries

Isolation of RNA from venom glands was performed as described previously [17]. Illumina sequencing libraries were generated using the TruSeq Stranded mRNA kit. The sequencing libraries were created using size selected RNA (300-800 bp). The venom gland libraries were sequenced in a single HiSeq2500 lane for 2 x 150 cycles producing paired reads 150 nucleotides in length.

Iso-seq libraries were generated using the SMRTbell Express Template Prep Kit 2.0 for the Sequel II system v8.0 with Sequel II Chemistry 2.0. Reads were processed using the Pacific

Biosciences recommended processing pipeline which included using the circular consensus sequence (CCS) reads to generate high fidelity (HiFi) reads. Next primers were removed from the HiFi reads to yield full-length (FL) reads. The FL reads were further refined through the removal of read concatemers to generate full-length non-concatemer (FLNC) reads. The FLNC reads were clustered to identify initial isoforms which were mapped (Minimap2) to the *C. atrox* reference genome [31]. Finally, the mapped and clustered isoforms were collapsed based on gene mapping to yield gene summarized isoform sets. This initial processing of reads contained transcripts that likely represented degraded RNAs or mis-mapped transcripts so further manually processing was done on the putative MP transcripts.

### Processing and analysis of venom gland RNA long read data

The alignments of the collapsed isoforms from the Pacific Biosciences (PB) pipeline to the metalloproteinase (MP) gene complex (∼1.2 mega base pairs and 30 annotated MP genes) were visually inspected using the Integrated Genome viewer (IGV) which identified many isoforms that aligned to different annotated MP genes but were incorrectly assigned (“collapsed”) to a single PB gene group. Additionally, many isoforms aligned to most of the coding exons for a gene but lacked sequence aligning to the start codon. Secreted venom proteins likely require a secretion signal so the isoforms lacking alignment to the start codon were removed from further analysis since they likely represent sequencing of degraded RNAs. These observations prompted us to refine the PB MP gene assignments by filtering the collapsed isoform set according to the following criteria: i) sequences that lack a start codon were removed ii) sequences that align within a gene interval bounded by 200 nt upstream of the start codon and 500 nt downstream of the stop codon were assigned to that gene. Filtering based on these criteria removed many sequences but retained a diverse set of putative isoforms from several classes (exon skipping, intron retention, truncation) (Fig S13 of *MDC4*). However, these curated sets of MP isoforms contained multiple full-length transcripts that were highly identical to each other. It is difficult to determine if these nearly identical transcripts represent biological differences (e.g. different untranslated region (UTR) lengths resulting from different transcriptional start sites (TSSs)) or artifacts of sequencing (e.g. degraded RNAs). Therefore, we specifically focused on accurate assessment of *MPO1* expression with general attention on MP expression variation between individuals. To do this we tailored our approach towards understanding quantitative between specimen expression differences of reference genome-derived full-length hypothetical MP transcripts.

### Analysis of venom gland gene expression using short and long read data

To quantify venom gland gene expression, we first constructed a venom gland meta-transcriptome that combined reference genome MP transcripts with sequences from venom gland PacBio isoseq for non-MP genes. This approach of combining annotated reference transcripts with unannotated isoseq reads represents a trade-off whereby we aim for accurate assessment expression of MPs at the gene-level while accepting the reduced accuracy of our non-MP expression differences. Next, reads from high throughput short read sequencing of venom gland RNA were pseudo-aligned (Kallisto version 0.48.0) to the meta-transcriptome [32]. Differential gene expression analysis was done using edgeR [33] and log_2_-transformed counts and log fold-change were visualized using R [34].

Our quantitative analysis of *MPO1* gene expression yielded a bimodal expression pattern with some individuals abundantly expressing *MPO1* while others had barely detected levels of *MPO1*. Thus, we sought to determine if specimens with low expression counts expressed a partial or full-length transcript at low levels with the assumption that partial transcripts are unlikely to make a functional MPO1 protein and therefore low *MPO1* expression could be interpreted as MPO1 absent from venom. To assess if low-expressing *MPO1* specimens expressed a full-length transcript at low levels or did not express a functional transcript at all, we needed to determine the limits of both sequencing technologies. First, to determine if full length *MPO1* transcript is detectable in the single molecule sequencing data of our non-reference specimens we inspected the processed putative isoforms that align to the reference genome. Two specimens (TX1 and TX3) have putative isoforms that align to the annotated *MPO1* exons with some potential isoform diversity (spliced exons, retained introns, novel exons) present in one of the non-reference specimens (Fig. S9; TX3). Whereas, in two specimens with relatively low *MPO1* expression we did not detect a full-length transcript (Fig. S9; AZ1 and TX4). This suggests that either in specimens with low *MPO1* expression a full-length transcript is not transcribed or it falls below the limit of detection for single molecule sequencing. The depth of short read sequencing did detect reads that mapped to the reference transcript (Fig. S7A) so to determine if those reads could yield a full length *MPO1* transcript we assembled venom gland transcriptomes for each specimen and aligned those assembled transcripts to the genome. We did not identify full-length assembled transcripts that aligned to all *MPO1* exons but for specimens with relatively high expression (NM1, TX3, TX1) we did observe partial transcripts that tiled across the genomic region and linked all coding exons (Fig. S8A). However, in the specimens with relatively low *MPO1* expression we found only a few partial transcripts linking several exons but did not identify a collection of partial transcripts that linked all exons (Fig. S8B).

We also mapped reads to the MP transcripts [35] and calculated coverage across the transcripts [36]. For *MPO1* we find, one region of the transcript, encoded by exons three and four, has zero coverage in all low expressing specimens but detectable low coverage in all high expressing specimens possibly indicating a difficult to sequence region or high sequence identity between exons three and four of MP genes reduces the number of unambiguously mapped reads (Fig. S7A, B). Therefore, any specimen with zero coverage only across this type of region (AZ3, Fig. S7A) may have low-level expression of a full-length transcript whereas specimens (AZ1, AZ2, TX4; Fig. S7A) with multiple zero coverage regions are less likely to express a full-length transcript. In summary, our analysis of MPO1 transcription suggests that the specimens with relatively low expression likely do not make full-length transcript. The detection of reads in specimens that are inferred to have deleted *MPO1* could be due technical or biological reasons. For example, rare index hopping on multiplexed sequencing runs could result in a very low level of reads from one specimen (*MPO1^+^*) being incorrectly assigned to another specimen (*MPO1-*). Incomplete deletion events during recombination events could remove most but not all exons or shuffle some exons to non-reference locations. If the product of sloppy recombination (partial genes) are near (or retain) active cis-regulatory elements then RNA may be detected even in the absence of a full-length gene.

We extended our analysis to *MDC8c* and for two specimens (Fig. 3, AZ2 and AZ3) with the lowest relative expression we identified a small set of partial transcripts that did not fully tile across all exons (Fig. S10A) whereas all other specimens had a full collection of tiling partial transcripts aligning to all exons and/or putative full-length isoforms (Fig. S10A, B). For *MAD3b*, the three specimens with the lowest relative expression also had a small set of partial transcripts that did not fully tile across all exons (Fig. S13A; AZ1, AZ2, AZ3) and for one of these specimens (AZ1) we also did not detect a putative full-length isoform (Fig. S13B) thus further supporting our position that when a venom gene has a bimodal expression pattern those specimens with relatively low expression likely do not make full-length transcript. The signature of bimodal expression appears to be key because only identifying specimens by the presence of a reduced set partial transcripts linking exons can be misleading. For example, in the case of *MAD3b* two specimens lacked a full tiling set of partial transcripts but did have putative full-length transcripts (Fig. S11 A, B; TX3 and TX4) and importantly, these two specimens expressed relatively high levels of *MAD3b* (Fig. 3A). For *MDC4,* we found in all specimens partial assembled transcripts linking all exons and a collection of putative full-length isoforms (Fig. S12A, B).

### Development of polyclonal antibodies with class-specific binding to venom metalloproteinases

To develop antibodies highly specific for individual MP proteins we first aligned the amino acid sequences of the MDC4 and MPO1 metalloproteinase domains and identified prospective peptide immunogens on the basis of predicted immunogenicity and sequence uniqueness. Candidate peptides (MDC4: 253-SNEDKITVKPEAGYT-267, MPO1: 358-RPGLTPGRSYEFSDDS-373) were selected, synthesized and coupled to a carrier before immunization of two rabbits (Pacific Immunology). To characterize the sensitivity and specificity of the antibodies, total *C. atrox* venom (0.5 micrograms per lane) was fractionated by SDS-PAGE and transferred to a PVDF membrane. After transfer the membrane was cut into strips corresponding to gel lanes and each strip was probed with pre- or post-immune sera at a range of dilutions. Sera from production bleed three was used for affinity purification of polyclonal antibodies using the immobilized peptide antigen.

To determine if the polyclonal-MDC4 antibody signal in a WB is specific for MDC4 we performed a competition experiment. We synthesized the orthologous antigen peptide sequence from the 14 class III MP paralogs, incubated peptide (10, 30, 100-fold molar excess) with MDC4-antibody (0.125 ug/ml) overnight with shaking at 4°C. The peptide-antibody or antibody only solutions were then used to probe WB membrane strips corresponding to a single gel lane loaded with one µg total *C. atrox* venom. The MDC paralog peptides do not reduce the band intensity suggesting our polyclonal antibody is specific for MDC4.

### Targeted sequencing of the Metalloproteinase genomic region

A hybridization-capture approach was used to enrich for genomic DNA encoding the metalloproteinase gene complex from individual specimens. Using the *C. atrox* MP reference sequence (∼1.2 mega base pairs (M bp) with 30 annotated MP genes) as a template, deoxyribonucleic acid (DNA) baits (100 nucleotides long) were designed and synthesized (Arbor Biosciences) at an average density of one bait every 125 nucleotides. However, an initial pilot experiment to evaluate hybridization efficiency and sequencing coverage across the MP complex (using genomic DNA from the same specimen used to generate the reference genome) identified genomic regions with low sequencing coverage. It was unclear if those regions were difficult to sequence or not present in the sequencing library because the absence of a bait resulted in poor enrichment. So additional baits were designed across the low coverage regions and then both the original and second batch of baits were used to enrich genomic DNA from all specimens. The Pacific Biosciences Multiplex Genomic DNA Target Capture Using SeqCap-EZ Libraries protocol was followed for generating the final sequencing libraries. Kapa HiFi polymerase was used for all amplification steps. The average length of input genomic DNA fragments for the sequencing libraries was seven kilo basepairs (kb) (range of two - ten kb). The mean DNA fragment length of the initial barcoded library was five kb and the mean fragment length of the final enriched DNA library was four kb. The libraries were pooled and sequenced with the PacBio Sequel II.

We first assessed if target enrichment long read sequencing isolated genomic sequence across the complete MP region by processing and analyzing reads from the specimen that was used previously to generate a bacterial artificial chromosome (BAC) library and a reference genome assembly [15,16]. Visual inspection (Integrated Genome Viewer (IGV), version 2.8.0) of reads aligned [37] (NGLMR version 0.2.7) to the reference assembly showed high coverage across the MP region but with high levels of variation. In this study we focused on the *MPO1* and *MDC4* genes and found complete cover across this region for the reference specimen (TX1; Fig. S18B) thus allowing us to conclude the absence of reads in this region for a particular specimen may reflect the absence of this genomic sequence in that individual.

### Assembly of targeted sequencing reads

Pacific Biosciences circular consensus sequence (CCS) reads were trimmed and corrected using CANU (version 1.9) [38–40]. The corrected reads were assembled using the Flye genome assembly program (version 2.8-b1674) [41]. The assembled contigs were aligned to the reference genome using LAST (lastal version 847) [35].

## Results

### Bimodal expression of several venom toxins in *C. atrox*

*C. atrox* venom is composed of metalloproteinases (MPs), PLA2s, serine proteinases, lectins, vasoactive peptides, L-amino acid oxidase and several other proteins consistently found in *Crotalus* species but at relatively low expression levels <u>[12,42]</u>. In order to capture the potential intraspecific variation in these components, we initially surveyed venoms from nine adult *C. atrox* specimens from across its range (Fig. 1). And in order to detect variation in protein composition at the level of resolution of single toxins, and to be able to differentiate among closely related toxin sequences, we performed mass spectrometry on whole venoms and used exclusive-unique peptide (EUP) counts of known venom proteins to estimate their relative abundance.

**Fig 1.**
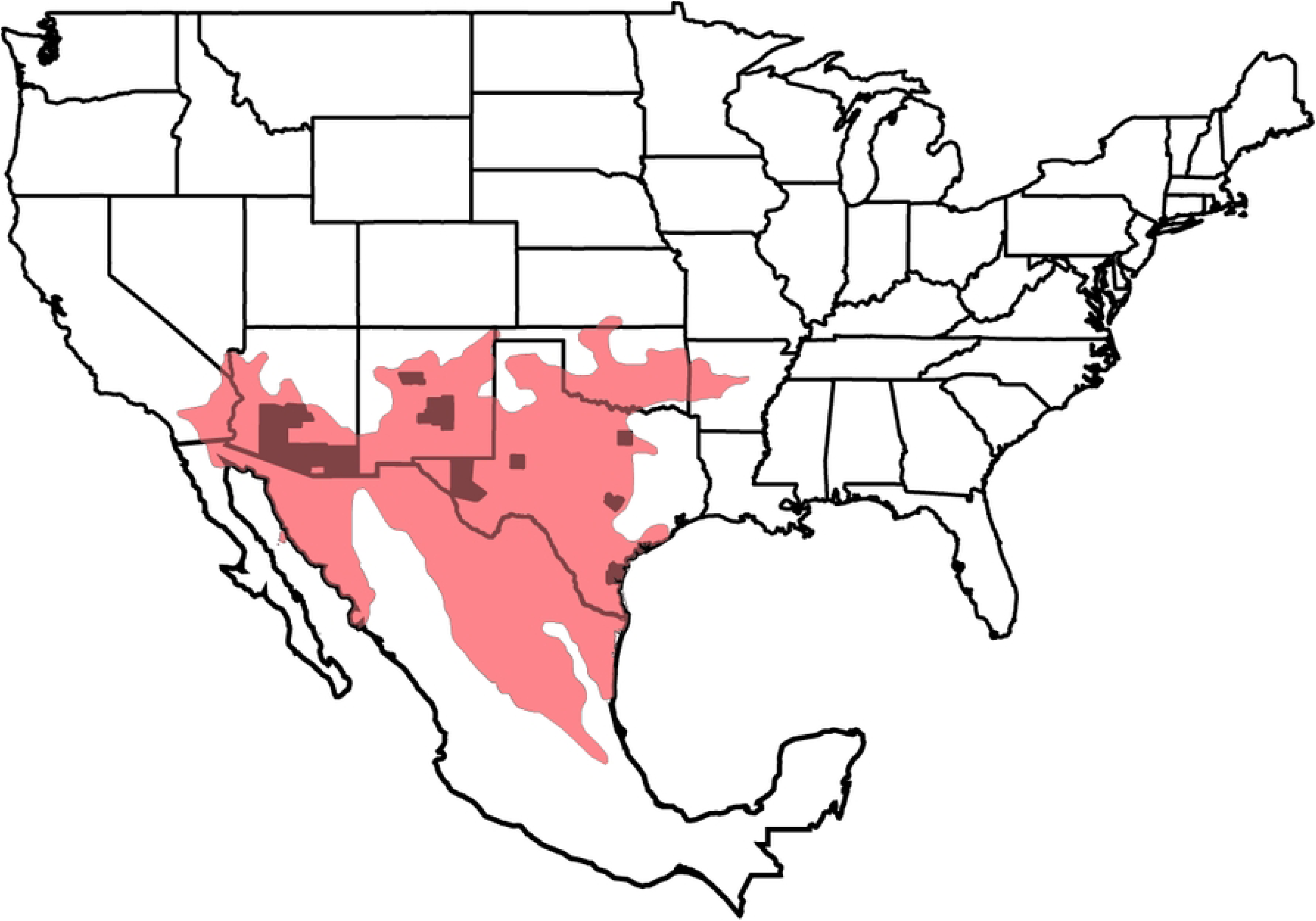
*C. atrox* species distribution and specimen sampling. The geographic distribution of *C. atrox* species (red shading) and specimen locations (black shading of counties) are shown.

Among the MPs, MDC4, MDC2, MDC7 and MAD3a exhibited limited variation in abundance (small spreads in EUP counts) between specimens. However, three MPs -- MPO1, MAD3b and MDC8c are relatively abundantly expressed in some individuals but are markedly reduced or lack expression in others (Fig. 2a). In contrast, most members of other venom protein families (e.g. phospholipases, serine proteinases) exhibit limited variation in abundance between individuals, except galactose-lectin (Fig. 2a, b and Fig. S1).

**Fig 2.**
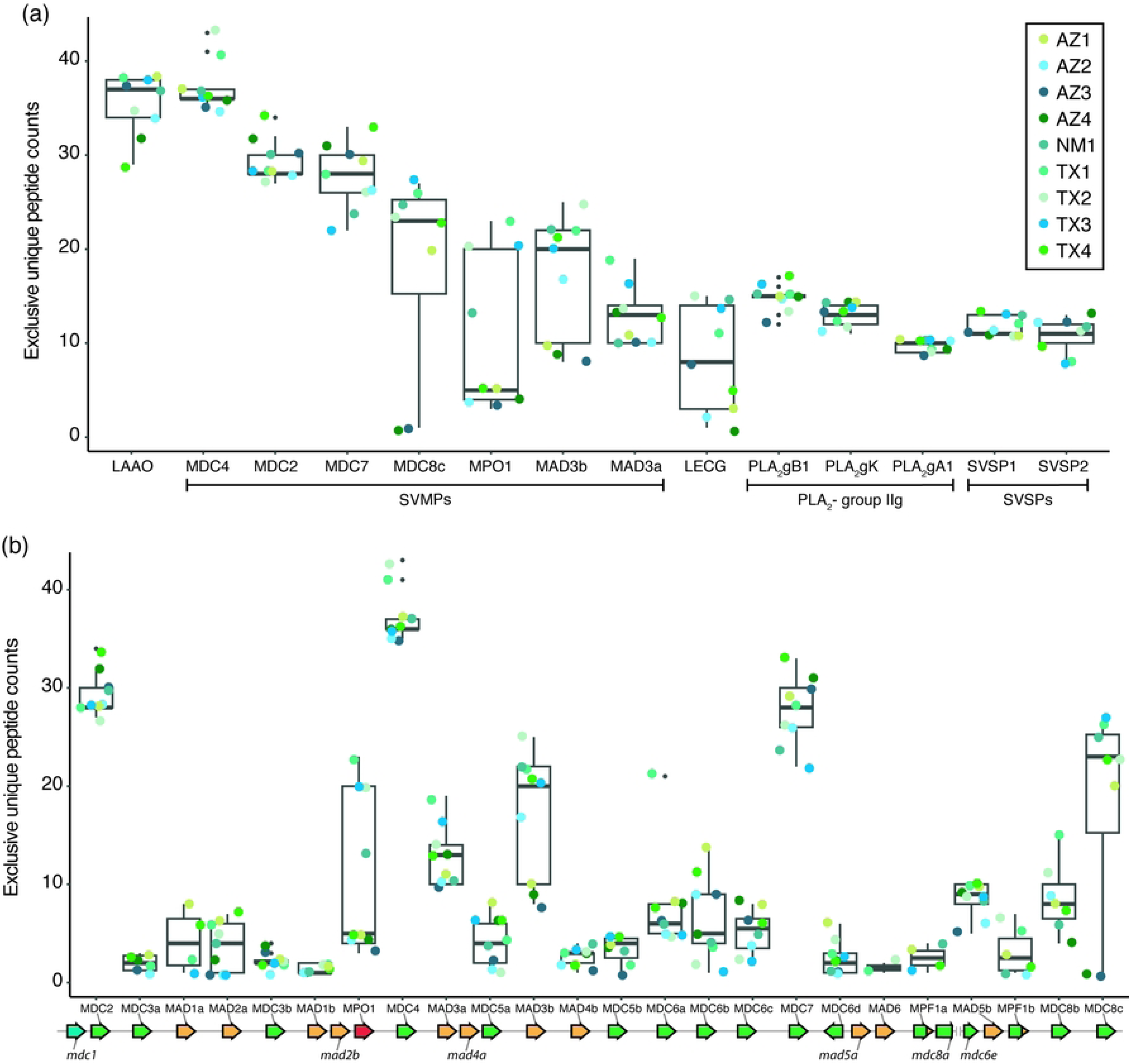
A bimodal expression pattern of several *C. atrox* venom toxins. (a) Average peptide counts for individual venom proteins from nine specimens (boxed legend in top right). Proteins are ordered along the x-axis from high to low abundance. Box plots show the upper and lower counts that span the interquartile range (IQR: 25 to 75th quartiles) and the median count (black line inside box). The extreme (1.5*IQR) lines extend from the top or bottom of the box and counts beyond the extremes are noted with small black dots. Most venom protein counts are similar between all specimens (narrow boxes) except MPO1, MAD3b, MDC8c and LECG counts vary widely between specimens (wide boxes). Venom proteins expressed at lower levels than than SVSP2 (Snake venom serine proteinase - 2) are shown in Supplemental Fig 1a. Average peptide counts for all known *C. atrox* metalloproteinases (MP). MPs are ordered along the x-axis according to the order of genes in the reference genome. The reference genome MP gene order is shown schematically below the box plots with genes represented as arrows (class III MPs in green, class II MPs in orange, class I MP in red and fusion MPs in green and orange). Expressed MPs have a line linking the protein identifier and gene arrow above the gene complex and undetected MPs have a line linking the respective protein identifier below the gene complex. The only unresolved gap in the reference sequence is also shown between *MDC8a* and *MDC6e*. MPO1, MAD3b and MDC8c have the broadest variation in expression between specimens. MDC2, MDC4 and MDC7 are the most abundant MPs and have limited variation in expression between specimens. Protein abbreviations: MDC, Metalloproteinase, Disintegrin, Cysteine-rich domains, class III zinc metalloproteinase; MAD, Metalloproteinase and Disintegrin domains, class II zinc metalloproteinase; MPO, Metalloproteinase domain only, class I zinc metalloproteinase; LAAO, L-amino acid oxidase; LECG, C-type lectin galactose, PLA_2_-gB1, Phospholipase A2, group IIG, B1; PLA_2_-gK, Phospholipase A2, group IIG, K; PLA_2_-gA1, Phospholipase A2, group IIG, A1.

To further ascertain the state of MP expression in certain individuals (low or absent), we determined that it was necessary to consider both the coverage and abundance of EUPs obtained by mass spectrometry. We aligned EUPs to hypothetical translations of MP reference amino acid sequences and calculated metalloproteinase domain coverage (Figs S2 - 5). This analysis reveals that peptides mapping within the metalloproteinase domain are detected in all specimens, however, the coverage for different segments of the protein (and abundance) varies significantly between specimens. For example, for MPO1, the coverage of peptides spanning the metalloproteinase domain is less than half (39 - 49 %) of the domain length in five specimens, whereas in four specimens most (> 90%) of the domain is covered (Fig. S2a and b). To determine if specimens expressing MPs at low levels are due to full coverage with low counts or partial coverage and low counts, we plotted the abundance (counts) of a subset of non-overlapping peptides that span the MP domain. This analysis reveals that the abundances of the few peptides found in the low-coverage specimens are extremely low relative to the abundances of the same peptide identified in the high coverage specimens (Fig. S2c). We also compared the mean counts for all MPO1 peptides detected in each specimen with MPO1 metalloproteinase domain coverage and found the specimens with low coverage also had very low mean counts (Fig. S3b, d). The positive correlation between low coverage and mean peptide counts is also found for the MDC8c (Fig. S3a - d) and MAD3b (Fig. S4a - d) proteins. Interestingly, we also note that the low abundance and coverage of MPO1, MAD3b and MDC8c are shared by three specimens (a fourth specimen has similar coverage and abundance only for MPO1 and MDC8c).

These observations are in striking contrast to, for example, those for the MDC4 protein, where the domain coverage and individual and mean peptide counts are very similar between specimens (Fig. S5a-d). The detection of just a few peptides for MPO1, MDC8c and MAD3b that only partially span the key protein domain raised the possibility that functional proteins are absent from these specimens’ venoms. This variation in toxin expression led us to investigate the potential genetic mechanism(s) that could be responsible.

### Lack of toxin protein expression is correlated with lack of toxin gene mRNA expression

Variation in toxin protein expression could be due to gene regulation (mRNA expression) or gene content. We sought to address whether regulation of venom gene transcription varies using two complementary sequencing platforms to qualitatively and quantitatively assess venom gland gene expression profiles of seven specimens. We used single molecule sequencing of full-length cDNAs from individual *C. atrox* venom gland RNA samples to generate representative venom gland transcriptomes that permits the qualitative analysis of both canonical and novel venom gland expressed transcript isoforms. Additionally, we used short read sequencing and mapping to MP reference transcripts to test if the observed variation for MPO1, MDC8C and MAD3b (Fig. 2) is consistent with mRNA transcript abundance.

We found that variation in MP mRNA expression broadly agrees with variability of venom protein abundance (Fig. 3a; genes as boxes and colored points represent individual specimens). For example, *MPO1, MDC8c and MAD3b* transcripts are abundantly expressed in individuals with relatively high venom peptide counts and coverage but reduced in the individuals that have low peptide counts and coverage (Fig. 3a). For example, specimens TX1, TX3 and NM1 express MPO1 near the level of the abundantly expressed MDC4 gene; but four other specimens (AZ1, AZ2, AZ3, TX4) exhibit reduced expression (Fig 3a). We observed similar bimodal patterns of *MDC8c* and *MAD3b* mRNA expression (stretched boxplots in Fig. 3a). The bimodal pattern of expression is not characteristic of all MPs since abundantly expressed class III MPs such as MDC2 and MDC7 are expressed at similar levels between specimens (narrow boxplots in Fig 3b). Furthermore, even close gene paralogs can have drastically different expression patterns (compare boxplots of MAD3a and MAD3b, Fig 3b) suggesting that MP gene expression may also be diverging.

**Fig 3.**
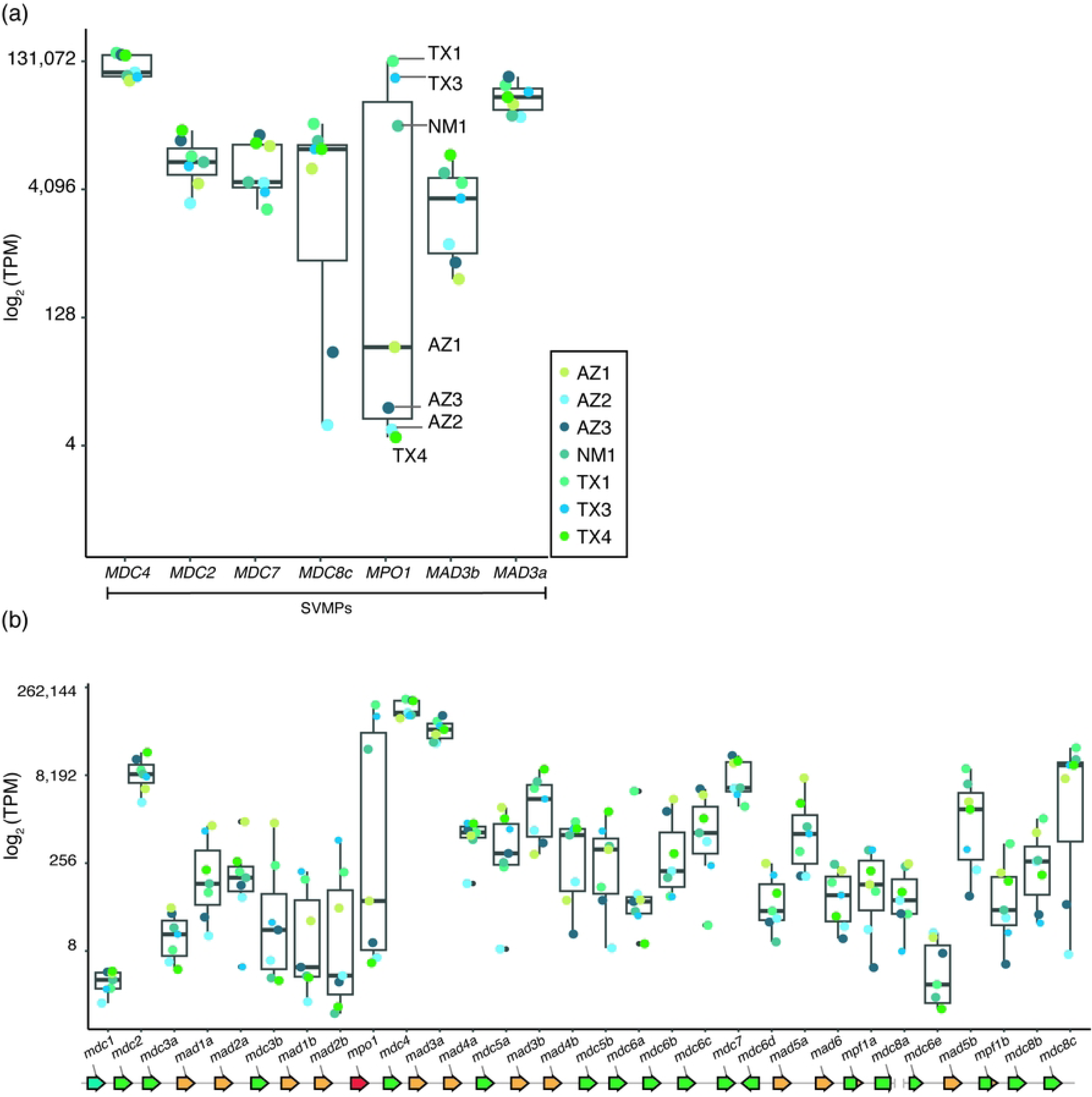
The bimodal pattern of toxin protein expression is also present in toxin RNA expression. Gene specific box plots of log_2_ transformed transcript counts (transcripts per million, TPM) for the main metalloproteinases (a) and all known metalloproteinases (b) presented in Fig 1a - b for seven individual specimens (boxed legend; colored points). *MPO1, MDC8c* and *MAD3b* have bimodal expression patterns (wide boxes) consistent with the protein expression. The expression pattern of most venom RNAs is consistent with the corresponding protein. The reference genome MP gene order is shown schematically below the box plots with genes represented as arrows as described in Fig 3.

To further investigate the nature of low MP transcript expression, we compared read counts (non-length normalized) of *MPO1*, *MDC8c*, *MAD3b* to the abundantly expressed *MDC4*. For MPO1, we found three specimens (TX1, TX3, NM1) with abundant mapped reads but four specimens with much fewer mapped reads (AZ1, AZ2, AZ3, TX4) (Fig. S6a, b). These specimens with fewer mapped reads also exhibited incomplete read coverage across the length of the reference transcript (Fig. S6c; AZ2 with ∼50%, AZ1 with ∼75%, AZ3 and TX4 with ∼85% coverage) with at least one zero coverage region (Fig. S7a, dashed boxes span zero coverage regions). This finding contrasts with specimens with high relative *MPO1* RNA expression (TX1, TX3, NM1) with read coverage several orders of magnitude greater across the transcript (Fig. S7b, compare the range of y-axis between A and B). This analysis suggests that specimens with relatively low *MPO1* expression likely do not make any full-length transcript (see Methods and Fig. S8,9).

Similarly, for *MDC8c,* the two specimens with the lowest relative expression (Fig 3, AZ2 and AZ3) had a small set of partial transcripts that did not fully tile across all exons (Fig. S10a), whereas all other specimens yielded a full array of tiling partial transcripts aligning to all exons and/or putative full-length isoforms (Fig. S10a, b). For *MAD3b*, the three specimens with the lowest relative expression also had a small set of partial transcripts that did not fully tile across all exons (Fig. S11a; AZ3, AZ2, AZ1) and for one of these specimens (AZ1) we also did not detect a putative full-length isoform (Fig. S11b). These results contrast with MP genes that have limited expression variability between specimens, for example *MDC4*, which shows a complete array of tiling partial transcripts, full-length isoforms and candidate variant isoforms (Fig. S12 - 13). We suggest that, as for *MPO1*, the specimens with relatively low *MDC8c* or *MAD3b* expression may not make any full-length RNA transcript.

### Design and characterization of specific antibodies to MPO1 and MDC4

We were particularly surprised to find MPO1 abundantly expressed in some individuals and apparently absent in others. MPO1 is the only P-I class MP in the species, whereas MAD3b and MDC8c both have at least one very closely related paralog and numerous more distantly related paralogs in *C. atrox*. To explore MPO1 expression and distribution in more specimens more easily, we endeavored to make polyclonal antibodies specific for MPO1 (and MDC4 – a low variation MP).

We inspected alignments of MP protein sequences to identify candidate peptide immunogens for the generation of specific polyclonal antibodies. One MDC4 peptide and two MPO1 peptides were synthesized and used to immunize rabbits. Reactivity to total venom in a Western blot assay was measured using crude sera and the sera with the highest activity were used as sources for the affinity purification of peptide-specific polyclonal antibodies. Titration of the affinity-purified polyclonal antibodies shows strong signal with minimal background (Fig. 4 a, b). The MPO1 antibody is highly specific to class I MPs because we observe a single band at approximately 25 kilodaltons (kDa), where the only known class I MP (MPO1) is predicted to migrate (Fig. 4b). Because several class III MPs are expressed in *C. atrox* venom (particularly MDC2, MDC7 and MDC8c) and those proteins are expected to migrate near MDC4, we needed to exclude the possibility that the signal from the anti-MDC4 antibody could be attributed to binding class III MPs other than MDC4. We performed a competition experiment by pre-incubating peptides from known class III MP paralogs with the anti-MDC4 antibody and examining their effect on the Western Blot signal (Fig. 4c). We find that while the MDC4 peptide competes for antibody binding and reduces the signal in a Western blot, the (paralogous) peptides from other class III MPs fail to reduce the signal (Fig. 4d).

**Fig 4.**
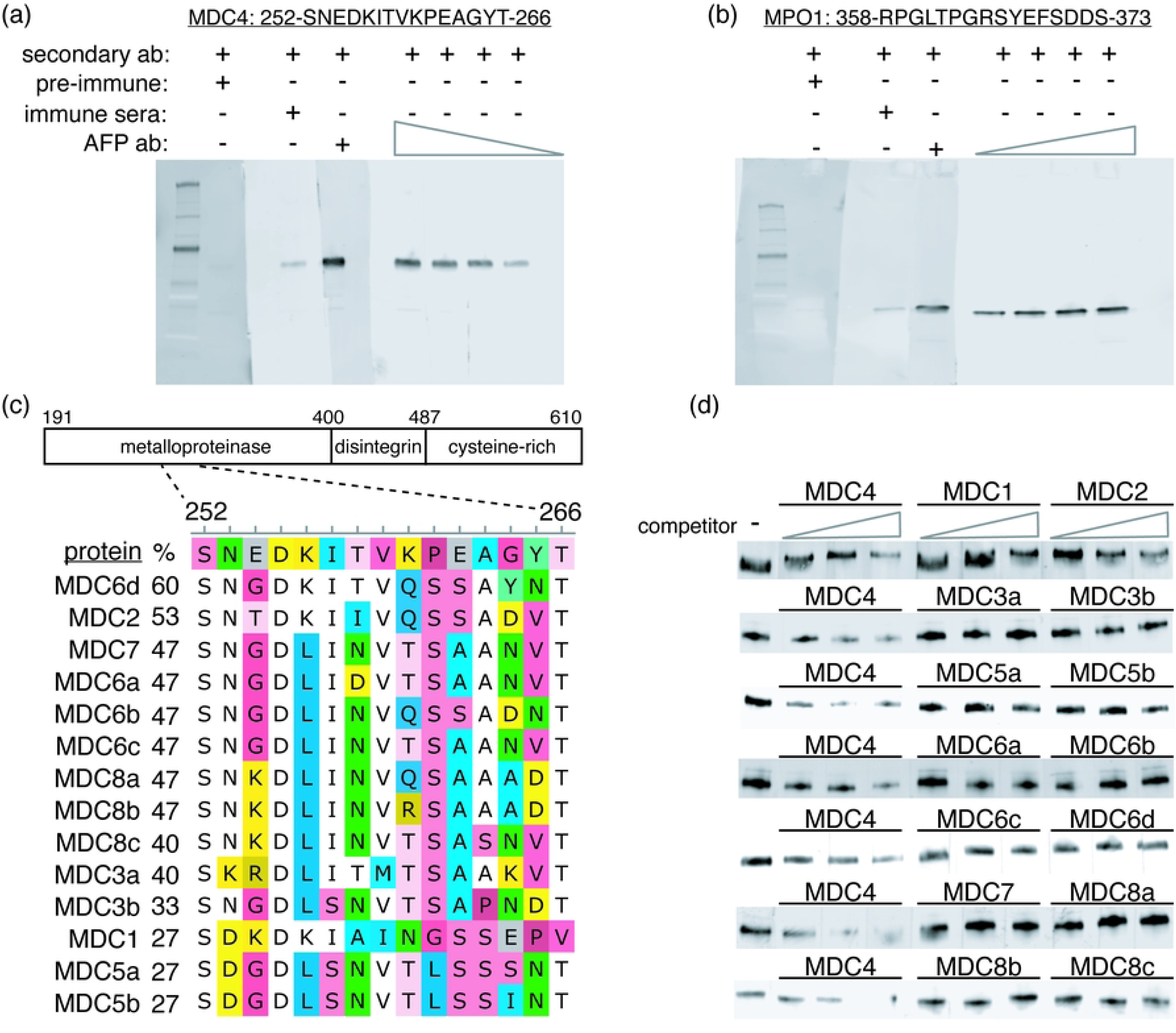
Development of specific antibodies to MDC4 and MPO1. The activities of the respective (a: MDC4, b: MPO1) affinity-purified antibodies (AFP ab) were titrated using total venom from the reference specimen (TX1) across a broad concentration range (1 to 0.125 micrograms per milliliter (µg/ml) using 2-fold steps). The sensitivity of both polyclonal antibodies is shown by the detection of respective proteins at the expected sizes (class III MP ∼48 k Da, class I MP ∼ 24 k Da) at increasing dilutions of antibody. The MP class specificity of each antibody is evident by the lack of bands at non-target sizes. To test the within-class specificity of the polyclonal-MDC4 antibody peptides corresponding to the orthologous sequences from MDC paralogs (c) were incubated with polyclonal MDC4 antibody (peptide molar excess of 10, 30 or 100-fold). The peptide-antibody mixture was then used to probe membrane strips containing transferred total venom. Polyclonal-MDC4 antibody mixed with MDC4 peptide has a titratable signal reduction whereas antibody mixed with peptide from MDC4 paralogs shows limited reduction of signal (d, compare signal with increasing competitor (three lanes below triangles) to lane without competitor (-)) suggesting the activity of polyclonal-MDC4 antibody has greater specificity for MDC4 relative to other MDC proteins that may be present in venom.

### MPO1 expression is low or undetectable in venoms from most *C. atrox* individuals in its western range

Our finding of variation in MPO1 abundance contrasts with a previous report of limited variation in *C. atrox* venom composition [43]. The difference in our results may be explained by the resolution of the methods employed, that is protein family-level resolution for SDS-PAGE versus single protein resolution when combining mass spectrometry with gene annotations. In addition, it is possible that our survey may have captured venom variation that could be linked to some variable, for example ecology or geography not represented or apparent in prior surveys (e.g. all of Rex and Mackessy (2019) specimens were from Arizona).

Indeed, we note that three of the four specimens that exhibited very low or no MPO1 protein or RNA expression (AZ1, AZ2, AZ3) originate from the western part of the species range while those exhibiting abundant MPO1 expression (TX1, TX3, NM1) originate from the eastern part of the range. Interestingly, Schield et al. (2015) previously identified a genomic signature of population structure in *C. atrox*. Specimens from each side (east or west) of the Continental divide (CD) were shown to have a distinct set of genetic markers. The observations of both venom variation and population structure in *C. atrox* raised the possibility that distinct venom compositions might occur in different populations.

To further investigate whether MPO1 expression correlates with geography, we examined 14 additional venom samples from across *C. atrox*’s range (along with the original seven samples) for the presence of MPO1 (and MDC4) using specific antibodies in a Western blot assay.

While *C. atrox* venom samples from across the range show similar levels of MDC4 expression (Fig. 5a - c), MPO1 is undetected (or relatively weak: AZ1, AZ5) in most (7/8, 88%) specimens found west of the CD (Fig. 5a). In contrast, for most eastern specimens, MPO1 is clearly detected (15/19, 79%) with four exceptions (Fig. 5b and c; undetected: NM7, TX4, TX6; relatively weak: TX5). We find statistical support for the association between geography and weak or undetectable MPO1 expression ((Χ^2^1, N=27) = 7.73, p=0.0054).

**Fig 5.**
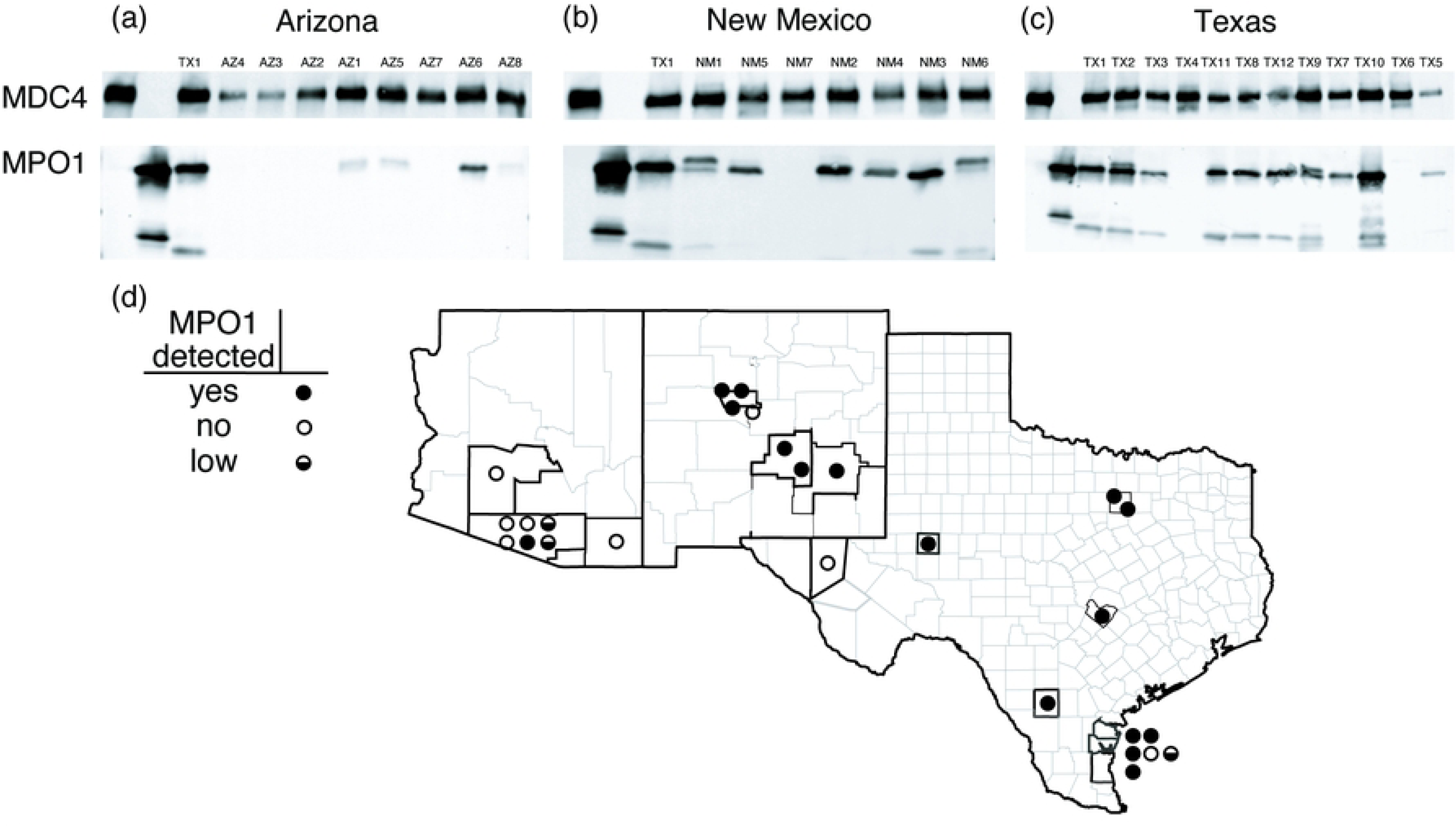
MPO1 expression is undetectable or low in venom from most western specimens. For each specimen (identifiers above the gel images) one microgram of total venom was separated by SDS-PAGE and transferred to a membrane. Additional controls on each gel are purified MDC4 (lane 1), purified MPO1 (lane 2) and total venom (lane 3) from the reference specimen (TX1; found east of the Continental divide). The top portion of each membrane was probed with anti-MDC4 and the bottom portion probed with anti-MPO1. MPO1 is undetected or extremely low (AZ1, AZ5, AZ8) in most (7/8, 88%) specimens west of Continental Divide (a, Arizona specimens). MPO1 is detected in most (15/19, 79%) eastern specimens (b, New Mexico; c, Texas) with several exceptions (NM7, TX4, TX5, TX6). In contrast, MDC4 is detected in all specimens with limited variability in signal. There is statistical support for an association between geographic origin (d; east or west of CD) and low or undetected MPO1 expression (7/8, west and 4/19, east; Chi-squared = 7.72, p-value = 0.005).

In our screening of many *C. atrox* venoms with the MPO1 polyclonal antibody, we also observed that in some specimens MPO1 appears as a doublet with a faster-migrating (lower molecular weight) band also detected. To test whether these bands are also the result of the polyclonal MPO1 antibody binding to the MPO1 epitope, we performed a competition experiment by incubating the polyclonal antibody with a molar excess of peptide and then probing membranes using the pre-incubated antibody. We find that both signals for the doublet and the lower molecular weight band are eliminated with increasing amounts of MPO1 peptide (Fig. S14), suggesting that our polyclonal antibody is detecting different forms of MPO1 in these samples.

The discovery of this correlation between MPO1 expression and geographical location prompted us to revisit our transcriptome data to perform differential gene expression (DGE) analysis between the east/west specimens as well from additional, independently collected specimens for which published venom gland transcriptomes are available [44]. The DGE analysis found that venom proteins with bimodal expression of protein levels (MPO1, MAD3b, MDC8c) also have a bimodal expression of transcript levels, however, differential gene expression is associated with geographical origin only for MPO1 (Fig. S15 - 16). This association raised the question of the potential genetic mechanism(s) underlying the absence of MPO1 expression in individual *C. atrox* animals.

### Whole gene deletion, a nonsense mutation and gene fusion within *MPO1* abolish MPO1 venom expression

To identify the genetic basis for loss of MPO1 mRNA and protein expression, we used targeted genomic sequencing, coverage analysis and annotation of assembled contigs from the *MPO1* genomic region of multiple individuals with both expression phenotypes (see Methods and Fig S17). We were able to identify mutations of various types that account for the low or no MPO1 expression phenotypes.

To validate our experimental approach, we first assessed coverage of the *MDC4* genomic region. We observe genomic sequencing coverage across the entire *MDC4* locus in all specimens which is consistent with an intact MDC4 locus driving full-length mRNA <u>expression</u> (Fig. 6a, filled boxes and Fig S18a, right half of all coverage plots). To assess the ability of our approach to recover the *MPO1* genomic region, we applied the method to the genomic DNA of the reference specimen (TX1) originally used to assemble the MP complex and found sequence coverage across the entire *MPO1* gene (Fig. 6a; TX1 and Fig S18a, b). Similarly, for one specimen with high MPO1 expression (TX3), we found sequence coverage across the MPO1 gene and subsequent annotation of assembled contigs yielded an intact *MPO1* gene that when hypothetically translated yields a full-length MPO1 protein (Fig. 6a and Fig. S18a).

**Fig 6.**
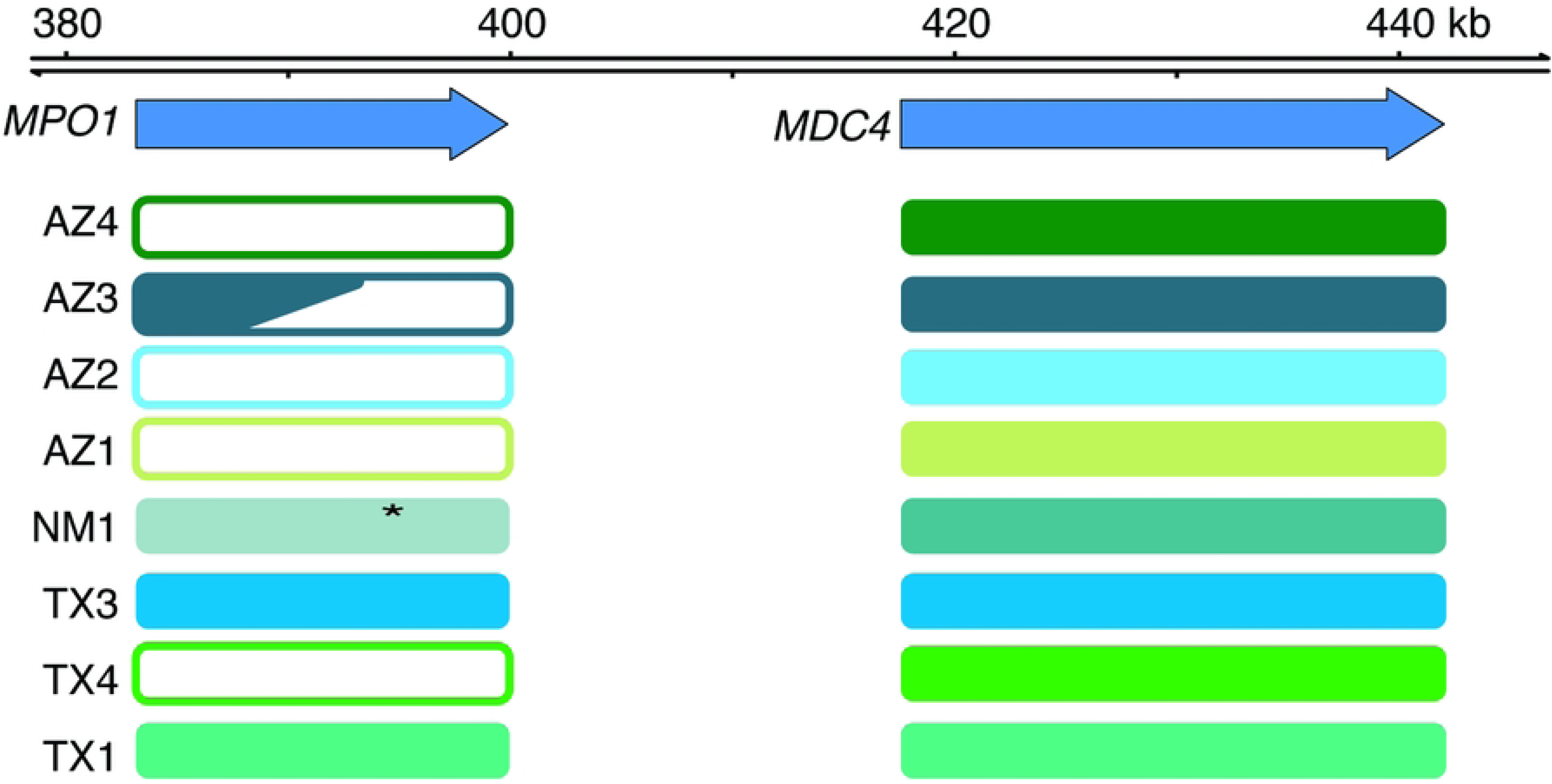
Whole gene deletion or mutation of *MPO1* exons underlies loss of MPO1 expression in venom. Schematic summary of aligned read coverage and annotation at the *MPO1* and *MDC4* loci. Four specimens (AZ1, AZ2, AZ4, TX4) have zero or low coverage at the *MPO1* gene but the adjacent *MDC4* gene has similar coverage relative to the other specimens. Annotation of assembled contigs from targeted sequencing reads identified one specimen (NM1) with a nonsense mutation in exon ten of the MPO1 metalloproteinase domain that is predicted to result in a truncated protein. Specimen AZ3 appears to have a broken *MPO1* locus resulting from a fusion event with *MAD5a*. A full-length intact *MPO1* gene is detected in specimen TX3.

However, annotation of another specimen (NM1) that yielded full coverage across the locus revealed that the *MPO1* gene contained nucleotide indels in exon ten that change the translation frame to a stop codon and, if expressed, would yield a truncated protein with only a partial metalloproteinase domain (Figs. S19a, b and S20c-f). These specimens with a mutated MPO1 are likely heterozygous for the mutation because our anti-MPO1 antibody detects a faint band co-migrating with full-length MPO1 protein (Fig. 5b).

In contrast, for three specimens (AZ2, AZ4, TX4) with undetectable MPO1 protein (Fig 5a, c) genomic coverage of the *MPO1* locus is not detected (Fig. 6a and Fig. S18a) and no assembled contigs are identified (Fig. S20a). We interpret these animals as bearing homozygous deletion mutations for the *MPO1* locus. We infer that the absence of MPO1 is an evolutionary loss, rather than a gain in eastern specimens, because previous work showed that the origin of MPO1 preceded the origin of the *Crotalus* clade [15].

Finally, annotation of another specimen (AZ3) revealed a chimeric *MPO1* gene with four exons (7 to 10) that perfectly match the amino acid sequence of MAD5a (Fig. S20b). Exon 11 matches both *MPO1* and *MAD5a* exon 11 with 63% similarity, but the following two exons match *MPO1* exons 12 and 14 with 98 and 100 % amino acid similarity (Fig. S20b). This suggests a recombination event between *MPO1* and *MAD5a* yielded a chimeric gene and the rearrangement of genomic sequence disrupted the gene encoding full-length MPO1.

In summary, we find in all specimens with low MPO1 expression evidence for *MPO1* inactivation via whole gene deletion, nucleotide insertion and deletion causing a frame shift and gene fusion. Gene inactivation by three distinct processes suggests the possibility that at least three independent loss of function events occurred in the gene encoding this toxin that is abundantly expressed when it is present.

## Discussion

We have shown that most individual venom toxins in the widely distributed rattlesnake *C. atrox* vary little in abundance between animals but that several toxins including three metalloproteinases have a bimodal pattern of variation -- expressed in some specimens but absent or barely detected in others. The variation in MPO1 expression is particularly surprising because, unlike MAD3b and MDC8c which have closely related paralogs expressed in *C. atrox* venom, MPO1 is the only toxin of its type (a P-I class metalloproteinase) in *C. atrox*, it is also one of the most abundant individual venom components (when present), and the absence of MPO1 expression occurs largely in animals found in the western region of the species’ range. Furthermore, we found that loss of MPO1 expression was not a consequence of transcriptional down-regulation but that several independent gene inactivating mutations have occurred (whole gene deletion, indel/frameshift, and a gene fusion) that eliminate the toxin from the venom arsenal. The repeated inactivation of MPO1 within a *C. atrox* subpopulation raises the questions of what ecological factors and evolutionary processes would cause a major toxin to be abandoned in part of the species range.

### Variation in prey may underlie variation in *C. atrox* MPO1 expression

The principal function of snake venoms is to subdue and kill prey. Venoms often contain dozens of different proteins that act singly or in concert to disrupt one or more critical physiological processes such as hemostasis or neuromuscular activity. One main action of *C. atrox* venom, like that of many other rattlesnakes and members of the family Crotalidae, is to disrupt hemostasis by destroying vascular integrity and interfering with blood coagulation. The major families of toxins responsible for these activities are the metalloproteinases and serine proteases which constitute up to 50% and 20% of the venom by weight, respectively, in *C. atrox* venom [42]. MPO1 is one of the most highly expressed MPs among the thirty venom MPs encoded in the *C. atrox* genome [15], It is also the only P-I class metalloproteinase present in the venom. Because this class of enzymes has been shown to have distinct activities on extracellular matrix and fibrinogen substrates, and a more diffuse distribution *in vivo* relative to P-II and P-III class enzymes [29], it seems likely that MPO1 contributes a distinct spectrum of activities to *C. atrox* venom. The absence of MPO1 expression in most animals in the western part of the species range and the abundance of the toxin in most animals in the eastern part of the range, coupled with evidence for ongoing widespread gene flow throughout the species entire distribution [45], may reflect that there are different requirements for venom function across *C. atrox*’s range. One prime candidate to explain this geographic variation in venom composition would be differences in the availability of susceptible prey.

A growing body of evidence suggests that snake venom composition co-evolves with the diversity, availability, and susceptibility of prey [9,46,47]. Among pit vipers, for example, a large-scale survey of venom composition and dietary breadth among many North American species (including *Crotalus*) found a positive correlation between greater venom complexity and the phylogenetic breadth of stomach contents [44]. And a recent study on the widely distributed *C. viridis* rattlesnake identified differences in venom composition along the north-south axis of the species’ geographical range that correlates with varying susceptibilities of different prey types [24]. In this case, an MP-rich and low myotoxin-expressing venom in snakes from the more southern region was associated with a more diverse diet consisting of mammals, reptiles, and birds, while snakes in more northern regions exhibited an myotoxin-rich, low-MP venom and preyed principally on mammals. This inversion in myotoxin abundance correlates with a potent lethal effect of the toxin on small mammals but a lack of effect on lizards [24]. The abundance of lizard prey declines dramatically in more northern latitudes, so venom composition in this case appears to be tailored to the availability of susceptible prey, not prey preference.

*C. atrox* is one of the largest and most widely distributed rattlesnakes, with one of the broadest habitat ranges, occurring across much of the southwestern United States and northern Mexico (Fig 4d). Numerous dietary studies have been conducted that reveal the species to be a generalist with a wide range of mammal and bird species observed in stomach contents, as well as lizards [48,49]. A comparative survey of specimens from across its entire geographic range did not reveal any significant differences in the classes of prey consumed (i.e. mammals and lizards) [50], so there is no general prey type that might explain an east-west difference in venom composition.

However, *C. atrox* in different regions may prey on different species of mammals. For example, while kangaroo rats (*Dipodomys* sp) are preyed upon throughout the species range [50], Texas animals have been reported to prey principally on woodrats (*Neotoma* sp.) and pocket mice (*Perognathus* sp.) [48]. And lagomorph consumption appears to be greater in western (Arizona) versus eastern (Oklahoma and Texas) specimens [48,51,52]. Regional differences in prey availability may influence venom composition, particularly if there are significant differences in prey susceptibility. Several mammalian species have been shown to be markedly resistant to *C. atrox* venom, such as the Mexican ground squirrel (LD_50_ is 13 times that of laboratory mice; [53]), the hispid cotton rat (LD_50_ is about 40 times that of laboratory mice; [54]), and the gray woodrat (LD_50_ is over 250 times that of laboratory mice; [54]). Moreover, resistance in some of these species has been shown to correlate with serum antihemorrhagic activity which is directed against venom metalloproteinases [54]. Thus, one possible factor underlying the loss of MPO1 expression may be the emergence of resistance to MPO1 and/or other MP activities in important prey.

In the absence of direct evidence for prey susceptibility shaping MPO1 distribution, however, we must remain open to other ecological or environmental factors that could vary across the species range and affect venom composition. For instance, polymorphism in *C. scutulatus* venom content was not linked to dietary differences, but to temperature and climatic variation which may influence venom composition more indirectly through, for example, foraging behavior or exposure to predators [22].

### Evolutionary loss of toxins: neutral and selective scenarios

In the ongoing co-evolutionary arms races between venomous predators and prey, the potential evolutionary advantages gained by the addition of new venom components or new toxin activities are obvious. Numerous investigators and studies have asserted that the composition of snake and other animal venoms is under strong selection, including balancing selection [55–57] but the evolutionary mechanisms governing the flip side of venom evolution – the loss of venom components, are not so clear. Assuming that the geographic distribution of the MPO1 toxin reflects different requirements for MPO1 and venom function across the range, two distinct evolutionary processes could explain the observed recurrent inactivation of the *MPO1* gene and absence of the toxin. First, if the source of selection that maintains MPO1 function in the eastern part of the range is absent in the western range, the relaxation of selection to maintain MPO1 function in the western population would allow for the recurrent inactivation of the *MPO1* gene and null alleles to persist under neutral evolution and genetic drift.

Alternatively, it is also possible that selection could favor the elimination of the toxin from venom. It is becoming better appreciated that there can be fitness costs to the production of traits, such that the loss of a trait may be favored when the source of selection for the maintenance of a trait is removed (reviewed in [58]). The production of venom incurs a metabolic cost, with one study reporting an 11% increase in *C. atrox* resting metabolic rates during periods of venom replenishment, which is 10-fold higher than the predicted rate for production of an identical mass of mixed body growth [59]. In the sea anemone as well, venom production entails a trade-off with growth and reproductive rates [11] and is repressed under stress [60]. Research on other secreted body fluids (mammalian milk, seminal fluids) also suggests that the production of these biologically important mixtures requires significant energy investment [61,62]. In the example examined here, it is plausible that individuals homozygous for *MPO1* null alleles that completely eliminate an ineffective and metabolically expensive toxin may possess a fitness advantage over those individuals expressing an ineffective toxin.

It is increasingly apparent that toxin gene loss occurs frequently in venomous snakes [16,17,63,64] and other venomous animals [57]. Evolution via gene loss has been proposed as a potentially important albeit less appreciated mechanism in adaptation to changing environments or novel ecological niches [65,66]. While gene loss can often be ascribed as a consequence, not a cause, of evolutionary adaptation or regressive evolution, it is also the case that testing for selection on loss-of-function/deletion alleles poses distinct experimental and analytical challenges [67]. Nevertheless, selection on loss of function alleles has been reported for a small subset of loci in a recent large-scale population genomic study of cavefish [68] as well as being inferred to contribute to adaptation in herbivores [69], cetaceans [70], fruit bats [71], nematodes [72], vampire bats [73], and mole rats [74]. While beyond the scope of this study, as further examples of toxin loss are documented, it will be valuable and necessary to obtain large-scale population genomic data to identify the potential evolutionary forces operating on such loss-of-function alleles.

## Availability of data and materials

The data underlying this study are available in the NCBI Sequence Read Archive under BioProject PRJNA1142048. Accession numbers for specific data (raw reads) for each specimen are available in tables on FigShare (https://doi.org/10.6084/m9.figshare.27244716and https://doi.org/10.6084/m9.figshare.27244740). Data tables associated with peptide counts (https://doi.org/10.6084/m9.figshare.26499178),

processed RNA (https://doi.org/10.6084/m9.figshare.26503990),

processed DNA reads (atx021: https://doi.org/10.6084/m9.figshare.27154119,

atx022: https://doi.org/10.6084/m9.figshare.27154110,

atx235: https://doi.org/10.6084/m9.figshare.27154098,

atx236: https://doi.org/10.6084/m9.figshare.27154095,

atx237: https://doi.org/10.6084/m9.figshare.27154083,

atx238: https://doi.org/10.6084/m9.figshare.27154080,

atx239: https://doi.org/10.6084/m9.figshare.27154068,

atx240: https://doi.org/10.6084/m9.figshare.27154059) and

assembled contigs (atx022: https://doi.org/10.6084/m9.Figshare.27146163,

atx235: https://doi.org/10.6084/m9.Figshare.27146163,

atx236: https://doi.org/10.6084/m9.figshare.27154005,

atx237: https://doi.org/10.6084/m9.figshare.27154023,

atx238: https://doi.org/10.6084/m9.figshare.27154026,

atx239: https://doi.org/10.6084/m9.figshare.27154032,

atx240: https://doi.org/10.6084/m9.Figshare.27154035) are available on FigShare.

## Authors’ contributions

N.L.D. designed research. N.L.D. and E.J.C. performed research. N.L.D. analyzed data. N.L.D. and S.B.C. wrote the paper.

## Acknowledgments

The authors thank Elda E. Sanchez and Mark Hockmuller at the National Natural Toxins Research Center, Texas A&M University-Kingsville for samples; Sara Goodwin from the Cold Spring Harbor Sequencing core for technical consultations; Jory van Thiel, Matt Giorgianni, and Rabindra Thakur for comments on the manuscript; and members of the Carroll lab for critical discussions related to this work.

## Supporting information

**S1 Fig. Average exclusive unique peptide counts for venom proteins.**

Average exclusive unique peptide counts for non-metalloproteinase venom proteins. Additional venom proteins are detected in the venom of most specimens at relatively low levels and vary little between specimens (Pla2g2g-C1, Phospholipase A2, group IIG, C1; SVSPC1, Snake venom serine proteinase-C1; SVSPH, Snake venom serine proteinase-H, TXVE, Snake venom vascular endothelial growth factor toxin; QPCT, Glutaminyl-peptide cyclotransferase; PRDX4, Peroxiredoxin-4; BKIP, Bradykinin inhibitor peptide; BNP, C-type natriuretic peptide).

**S2 - S5** Figs. Individual peptide locations, coverage and counts for single proteins.

These figures show the individual peptide locations (A), percentage of protein coverage (B), individual peptide counts (C) and average peptide counts of a protein (D) for MPO1 (Figure S2), MDC8c (Figure S3), MAD3b (Figure S4) and MDC4 (Figure S5). For figures S2 - S5 (A), the amino acid locations (x-axis) of exclusive unique peptides (barbell shaped line segments) are mapped to a linear representation of the respective venom protein. Each row shows peptides from a single specimen with the specimen identifier on the right side of the plot and aligned with the horizontal bar plot (B) showing observed coverage across the metalloproteinase domain. Observed coverage is the percentage of the metalloproteinase domain covered by unique peptides after removal of conserved (non-unique between paralogs) or not detected sequences. A few sequence segments of the metalloproteinases are highly conserved among MPs so peptides from those regions cannot be unambiguously assigned to single proteins and have been removed from this analysis (blank spaces shared by all specimens). The counts (x-axis) of individual non-overlapping peptides (y-axis) spanning the protein are shown for each specimen (colored dots) (C). This analysis shows that when a protein has high coverage comprised of many peptides then the associated counts of those individual peptides is often uniform and consistent with the mean count for the total protein (D).

**S6** Fig. Short read counts and coverage of mapped reads to four reference transcripts (*MAD3b, MDC4, MDC8c, MPO1)*.

(A) Box plots showing between specimen variation in counts of mapped reads to reference transcripts. These counts are not length-normalized so longer transcripts can have higher counts and comparisons across the different classes of transcripts are not indicative of expression differences.

(B) Zoomed view of the MPO1 counts highlighting the variation between specimens.

(C) Horizontal bar plots showing the percent of the reference transcript covered (x-axis) by mapped reads from each specimen.

**S7** Fig. Read coverage across the *MPO1* reference transcript.

Abundant *MPO1* expression correlates with complete read coverage across the *MPO1* transcript.

Specimens with low total read coverage (A) have gaps in coverage (dashed boxes) across the *MPO1* transcript whereas specimens with high *MPO1* expression (B) have complete coverage across the transcript.

**S8** Fig. Assembled transcripts aligning to the *MPO1* gene and linking multiple exons Differences in completeness of assembled transcripts correlates with *MPO1* expression level.

(A) Specimens with relatively high *MPO1* expression have assembled transcripts that tile across the complete gene and link exons. (B) Specimens with relatively low *MPO1* expression lack assembled transcripts that tile across the complete gene and link exons. Additionally, specimen NM1 has a linking transcript that connects exon 8 to exon 11 and lacks alignments to exons 9 and 10 thus suggesting the mutated exons 9 and 10 may be removed through splicing.

**S9 Fig.** Detection of full-length and novel *MPO1* isoforms with single molecule sequencing. Detection of full-length and novel *MPO1* isoforms using single molecule sequencing. For two specimens (TX1 and TX3) a full-length *MPO1* isoform is detected. For specimen TX3, novel isoforms consisting of spliced exons, novel exons and/or retained introns are also shown. For two specimens (AZ1 and TX4) with low MPO1 protein and mRNA levels no full-length *MPO1* isoforms are detected.

**S10 Fig.** Assembled transcripts aligning to the *MDC8c* gene and linking multiple exons Consistency between assembled transcripts linking all exons and single molecule sequencing of full-length *MDC8c* isoforms. (A) *MDC8c* assembled transcripts that tile across the complete gene and link exons with the notable exceptions of AZ2 and AZ3. (B) Full-length *MDC8c* isoforms identified using single molecule sequencing.

**S11 Fig.** Assembled transcripts aligning to the *MAD3b* gene and linking multiple exons Single molecule sequencing recovers full-length *MAD3b* isoforms in specimens that lack assembled transcripts tiling across the gene. (A) *MAD3b* assembled transcripts that tile across the complete gene and link exons with the exceptions of AZ1, AZ2, AZ3, TX3, TX4. B) Full-length *MAD3b* isoforms identified using single molecule sequencing with the exception of AZ1.

**S12 Fig.** Assembled transcripts aligning to the *MDC4* gene and linking multiple exons Evidence for expression of full-length *MDC4* expression in all specimens.

(A) *MDC4* assembled transcripts from all specimens tile across the complete gene and link all exons. (B) Full-length *MDC4* isoforms identified using single molecule sequencing.

**S13 Fig.** Isoforms of *MDC4* are detected but often expression is limited to a subset of individual specimens.

(A) Isoform sequencing using Pacific Biosciences technology (IsoSeq) from four specimens (TX1, TX3, TX4, AZ1) followed by quantification of isoform abundance using short reads from all specimens identified isoforms from several general classes (grey boxes on left showing full-length, splicing, truncation) that align to the genomic region of *MDC4* (blue rectangles or arrows show regions of isoform sequence that align, green rectangles in top row show location of annotated *MDC4* exons. (B) Flipped box plots with the isoform-level counts (Transcripts Per Million) for seven specimens. Expression is limited or not detected in most specimens for most isoforms. For the three putative full-length isoforms (PB.1752.158_atx240 (TX4), PB.5518.684_atx239 (TX3), PB-3656.442_atx238 (AZ1)) hypothetical translations of only one (PB.1752.158_atx240 (TX4)) yields a full-length protein with the other two isoforms containing nucleotide substitutions that yield a hypothetical truncated protein.

**S14** Fig. MPO1 reactive doublet is competed away with MPO1 peptide.

Signal from “MPO1 doublet” and lower bands are reduced with excess MPO1 peptide. Initial characterization of the polyclonal MPO1 antibody and screening of individual *C. atrox* venoms revealed a doublet at ∼25 k Da and a lower molecular weight band. These bands may represent non-specific binding activity of the antibody, recognition of post-translationally modified MPO1, an MPO1 allele (doublet) and/or proteolytic processing of MPO1 (lower band). To distinguish between these possibilities we synthesized the MPO1 peptide antigen and mixed with the polyclonal MPO1 antibody (10, 30, 100 molar excess peptide). These MPO1-antibody-peptide mixtures were used to probe membrane strips from individual venoms (TX1, TX2, TX9, NM1, NM6). (E) The band intensities of the doublet and lower molecular weight decreased with increasing peptide supporting the possibility that the polyclonal antibody is detecting non-canonical MPO1 products (relatively fast and slow migrating polypeptides) that may represent post-translationally modified or proteolytically processed forms of MPO1.

**S15 Fig.** Differential gene expression between *C. atrox* specimens from east and west of the Continental Divide.

*MPO1* expression correlates with geographic origin with high *MPO1* expression in specimens found east of the Continental Divide (CD) and low expression in specimens found west of the CD. This motivated us to revisit our venom gland transcriptomes and perform differential gene expression analysis on east/west venom gland transcriptomes. We identified several venom metalloproteinase genes that are differentially expressed (A, B; blue fill of ball-and-stick, P < 0.001) between the east and west groups. The x-axis shows direction of fold changes (log-transformed; logFC) as a ball-and-stick pointing upwards (higher expression in west vs east) or downwards (lower expression in the west vs east).

In addition to *MPO1*, we also identified four additional MP genes (*MAD3b, MAD4b, MDC6b* and *MDC6e*) that are differentially expressed (supplementary fig. S15a and b). We included *MPOC* (a class I MP variant described in previous work but absent from our reference genome) in our analysis and found this gene to also be differentially expressed between east and western groups (supplementary fig. S15a and b).

**S16 Fig.** Independent support for the differential expression of *MPO1* in *C. atrox* populations.

Reads from published *C. atrox* venom gland trancriptomes (Holding 2021, PNAS) that were annotated as east or west are used with the same gene set in supplementary figure S15 to determine if any venom metalloproteinase genes are differentially expressed between the east/west groups. The variance in expression levels for all known venom metalloproteinase genes (A) for ten specimens (7, east; 3, west) generally follows the pattern shown in supplementary figure S15a. The same DGE analysis using the published venom gland data, reveals similar decreased *MPO1* expression (pink highlight, supplementary fig. S16a; P < 0.01) As in Figure S15 ball-and-stick plots below the respective box plots show the log fold change in expression for specific genes between the east/west groups. Filled blue circles highlight differentially expressed genes between the east and west groups (P < 0.01 in B).

**S17 Fig.** Targeted sequencing nucleic acid probe density and locations (black bars) at the *MPO1, MDC4, MAD3a* regions.

Across the entire MP gene complex probes were at an average density of one probe every 125 nucleotides but there are some larger gaps due to low sequence complexity.

**S18 Fig.** Low read coverage suggests the absence of *MPO1* in most western specimens.

(A) Aligned read coverage at the *MPO1* and *MDC4* loci for four western (AZ1 - 4) and eastern (TX1 - 3 and NM1) specimens. Four specimens (AZ1, AZ2, AZ4, TX4) have zero or low coverage at the *MPO1* gene but the adjacent *MDC4* gene has similar coverage relative to the other specimens. Prior work generated a reference genome using specimen TX1 and targeted genomic sequencing of TX1 yielded high cover across the total *MPO1* - *MDC4* region (upward arrows indicate coverage exceeds the y-axis). (B) High genomic read coverage at the *MPO1- MDC4* region for two specimens (TX1 and AZ2) with very high coverage.

**S19 Fig.** Targeted genomic sequencing of NM1 yields high coverage across the *MPO1* gene and reveals putative nucleotide mutations.

(A) Alignment of NM1 reads to the reference genome shows high coverage across the *MPO1* gene with high levels of nucleotide variation. Below the coverage histogram is a pileup of the individual aligned reads (horizontal grey strips) with each nucleotide that differs from the reference sequence shown as a colored tick mark. Zooming in on exons 10 (B), 11 (C) and 14 (D) shows the nucleotide substitutions and indels (downward spikes in coverage histogram and white gaps with horizontal bar in the read pileup). NM1 exon 14 has nucleotide variation but no indels are detected and a hypothetical translation of exon 14 is 100 % identical to the reference sequence.

**S20 Fig.** Alignment and annotation of *MPO1* and *MDC4* contigs supports inference of genomic deletions at the *MPO1* region.

(A) Contigs that align to the *MPO1* region were annotated to identify exons. For specimen AZ3, contig 37 aligns to the 5’ region of *MPO1* and contains intact exons 1 to 6 which encode the signal peptide and pro-domain. Contig 5 aligns to *MDC4* and extends towards *MPO1* but aligns poorly (dashed region). (B) Annotation of this sequence revealed metalloproteinase exons 7 through 12, plus 14. Amino acid percent similarity of exons 7 - 10 suggests these are *MAD5a* exons but exons 12 and 14 are more similar to *MPO1* (*MAD5a* does not contain an exon 14) thus this may be a chimeric gene fusion created during a recombination event. In the reference assembly *MAD5a* is not adjacent to *MDC4*. Annotation of the aligned contigs from NM1 and TX3 generally supports the identification of *MPO1* on these contigs. However, annotation exon 10 in NM1 reveals indels (compare C and E alignments at positions 13, 14 and 41) that is predicted to result in translation stop (position 24 in D and lack of similarity to reference (TX1) sequence). (F) Accounting for the indels in a hypothetical translation shows the amino acid sequence similarity between NM1 and TX1 exon 10 sequences.

